# Patterns of smallpox mortality in London, England, over three centuries

**DOI:** 10.1101/771220

**Authors:** Olga Krylova, David J.D. Earn

## Abstract

Smallpox is unique among infectious diseases in the degree to which it devasted human populations, its long history of control interventions, and the fact that it has been successfully eradicated. Mortality from smallpox in London, England, was carefully documented, weekly, for nearly 300 years, providing a rare and valuable source for the study of ecology and evolution of infectious disease. We describe and analyze smallpox mortality in London from 1664 to 1930. We digitized the weekly records published in the London Bills of Mortality and the Registrar General’s Weekly Returns. We annotated the resulting time series with a sequence of historical events that appear to have influenced smallpox dynamics in London. We present a spectral analysis that reveals how periodicities in smallpox dynamics changed over decades and centuries, and how these changes were related to control interventions and public health policy changes. We also examine how the seasonality of smallpox epidemics changed from the 17th to 20th centuries in London.

## Introduction

Smallpox was declared eradicated 40 years ago [1], after unparalleled devastation of human populations for many centuries [2,3]. Until the 19th century, smallpox is thought to have accounted for more deaths than any other single infectious disease, even bubonic plague and cholera [2–5]. In London, England, alone more than 320,000 people are recorded to have died from smallpox since 1664.

Investigation of smallpox dynamics in the past is important not only for understanding the epidemiology of a disease that has been exceptionally important in human history, but also in the context of the potential for its use as a bioterrorist agent in the future [6–10]. From the historical perspective, key issues include relationships between patterns of smallpox outbreaks and demographic changes, uptake of preventative measures, and interactions between epidemics and other historical events.

We examined 267 years of weekly smallpox mortality records from the city of London, England, beginning in 1664. The data span an early era before any public health practices were in place, the introduction of variolation and then vaccination, and then the decline of smallpox mortality until it became an extremely unusual cause of death. In addition to a statistical description of the temporal patterns in the data, we present a timeline of major historical events that occurred during the epoch we studied. Overlaying the historical timeline with smallpox mortality and prevention patterns provides an illuminating view of three centuries of smallpox history.

Some previous work on smallpox dynamics in London has been based on annual records [11,12], which lack the temporal resolution of the data we study, and consequently smoothed out seasonal patterns that we identify. More recent work has examined individual death records in a single London parish over a period of several decades (1752–1805) and identified a decline in adult smallpox mortality risk after 1770 [13].

To provide context, we briefly review the known history of smallpox, and describe its natural history, before presenting and analyzing the data on which we focus.

### Smallpox history

The origin of smallpox is not known. It appears to have coexisted with humans for thousands of years, and is thought to have first appeared in Asia or Africa sometime after 10,000 BC before spreading to India and China [2]. The earliest credible evidence of smallpox existence in the ancient world comes from the Egyptian mummies of the Eighteenth and Twentieth Dynasties (1570 to 1085 BC) [1,2].

After having established in Egypt, India and China, smallpox spread to Athens and Persia. By the eighth century AD the virus had reached Japan in the East and Europe in the West [2]. In the fifteenth century smallpox was widespread throughout Europe [1]. The Spanish invasion of Mexico in the sixteenth century brought smallpox to previously unexposed populations of the Aztec and Inca civilizations. Devastating epidemics killed nearly half of the native population of Mexico in less than six months after its first appearance in April of 1520 [2]. Two centuries later, smallpox had become a major endemic disease everywhere in the world.

Smallpox was directly responsible for the death of hundreds of thousands of people each year during the Early Modern period (1500-1800) in Europe [3]. In many cases surviving victims were left blinded or disfigured for life [14]. Smallpox also could have been a major cause of infertility in male survivors [2,3]. The only defence against this “speckled monster” was a procedure initially called *inoculation*, and now known by the more specialized term *variolation*. It was introduced in Europe only in 1721 despite being successfully used in China and India for centuries before. It can be described as an injection of the smallpox virus taken from a pustule or dried scabs of a person suffering from smallpox into a healthy individual [3]. The discovery of *vaccination* by Edward Jenner in 1796 provided a safer and much cheaper alternative to variolation. The original form of vaccination was an injection with cowpox virus, which also provided immunity to smallpox. This discovery was a major milestone in modern medicine as it laid the ground for all future vaccination strategies. Indeed, the word *“vaccine”* takes its origin from the latin *“vacca”* meaning cow, and was first used by Jenner to describe his new method of “vaccine inoculation” [15].

The discovery of a vaccine was the key factor that made eradication of smallpox an achievable goal. Other important factors were the absence of an animal reservoir, easy recognition of the disease from its symptoms, and non-existence of asymptomatic cases [16]. The World Health Organization launched its eradication campaign in 1967 [1,17]. Ten years later the world’s last endemic smallpox case was registered in Somalia. In 1980, a Global Commission declared smallpox eradicated [18]. This was the first disease to be eradicated entirely by human efforts. The only remaining viral samples were stored in laboratories in Russia and the United States [9]. “The greatest killer” [2] has never circulated since.

Unfortunately, the tragic events of September 11, 2001 and the anthrax scare of 2001 led to widespread concern about bioterrorism and, in particular, the use of smallpox as a bioweapon. In fact, its high case fatality rate, person-to-person transmission, and limited immunity in the population make smallpox even more dangerous than anthrax. The US Centres for Disease Control and Prevention (CDC) classifies smallpox as a Category A bioweapon agent [10,19].

An increasing proportion of the world’s population lacks immunity to smallpox [20,21]. As routine vaccination was stopped in the 1970s, most people born since then are fully susceptible. In addition, the level of immunity in people vaccinated before 1970 is unknown. Naturally acquired smallpox and variolation typically produce life-long immunity. In contrast, the *vaccinia* virus, which is the basis of the modern vaccine, provides complete immunity only for a short period of time, 3-5 years, after which immunity wanes rapidly [1,22]. Long term vaccine-induced immunity is uncertain. However, even after 20 years the case fatality rate in vaccinated persons is much smaller than in the unvaccinated [1].

The smallpox vaccine that was used until eradication is not considered safe, because of rare—but extremely dangerous—complications that include severe eczema, postvaccinal encephalitis (inflammation of the brain), and significant risk of death. During routine vaccination in the United States six to eight children died each year because of various vaccine-induced complications [2, p. 294]. Moreover, smallpox vaccination was not safe for people with weakened immune systems (e.g., pregnant women, HIV-positive individuals and cancer patients). Consequently, mass vaccination was ceased because the risks of complications were perceived to be much greater than the risk of a bioterrorist attack. In recent years, new and safer vaccines have been developed [23,24], which would likely be more pallatable to the population if mass vaccination were reintroduced.

Until 2018, vaccination was the only available measure to protect against smallpox. In July 2018, the U.S. Food and Drug Administration (FDA) approved a new antiviral drug (*tecovirimat*) as the first drug intended to treat smallpox [25]. However, the approval was based only on animal trials, as it was not feasible to conduct efficacy trials in humans [10,26]. Therefore the actual efficacy of this drug in humans is unknown.

### Types of smallpox

Smallpox is an acute, highly contagious and frequently fatal disease. The commonly accepted term “small-pox” was first used in England at the end of the 15th century to distinguish it from syphilis, which was known as “great-pox” [2, pp. 22–29].

Smallpox is characterized by high fever and a distinctive skin rash that often leaves pock scars after scabs fall off [14]. Infection usually occurs via the respiratory tract through air droplets if exposure occurs from face-to-face contact with an infectious person. Direct contact with virulent rash, bodily fluids, or bedding, blankets, or clothing used by an infectious individual can also, in rare cases, result in smallpox infection [2, p. 3] [1, pp. 121—168].

Smallpox is caused by the *Variola* virus, which takes its name from the latin word *varius* meaning spotted or *varus* meaning pimple. *Variola* is a member of the genus of orthopoxviruses, which also includes cowpox, monkeypox, buffalopox, vaccinia and many other members [2, p.6] [1, pp. 69-120]. There are two distinct variants of *Variola* that can cause smallpox: *Variola major* and *Variola minor* [1, pp. 1–68] [2, pp. 3–9] [22, pp. 525–527]. These two smallpox variants differ substantially in severity of symptoms and case fatality proportion.

*Variola major*, which had a case fatality of 5–25% and occasionally higher [1, p. 4], was the only known smallpox type until the beginning of the 20th century. Based on strain virulence and host response, a number of clinical types of *V. major* were defined [1, pp. 1–121]:

- **ordinary-type** was the most common (about 80% of cases), with case fatality 20%;
- **modified-type** was milder, had an accelerated course of infection, and occurred mostly in previously vaccinated persons;
- **variola sine eruption** was rare, occurred in vaccinated persons through contact with smallpox patients, and was characterized by a high fever without rash eruption;
- **flat-type** was characterized by lesions that remained flat. It was rare and usually fatal;
- **haemorrhagic-type** is distinguished from other types by especially severe symptoms, a short incubation period, and high case fatality ~ 96%. It was rare, occurred mostly in adults, and at the early stage of infection was hard to distinguish from ordinary-type or modified-type.

*Variola minor*, first recognized in 1904 [2], caused a milder, less virulent form of smallpox and had a mortality rate of about 1% or less. *Variola minor* was the only endemic type of smallpox present in England after 1920 [2, pp. 8,97]; [1, pp. 243]. In 1935, England became smallpox-free for the first time in history, and further cases of naturally acquired smallpox arose only from importation [2].

### Natural history of smallpox infection

The course of a single smallpox infection (its natural history) depends on the variant of the virus, the clinical type, and the vaccination status of the host individual. Since the ordinary-type of *Variola major* was the most common type of smallpox, we describe its natural history here (Fig 1).

**Fig 1.**
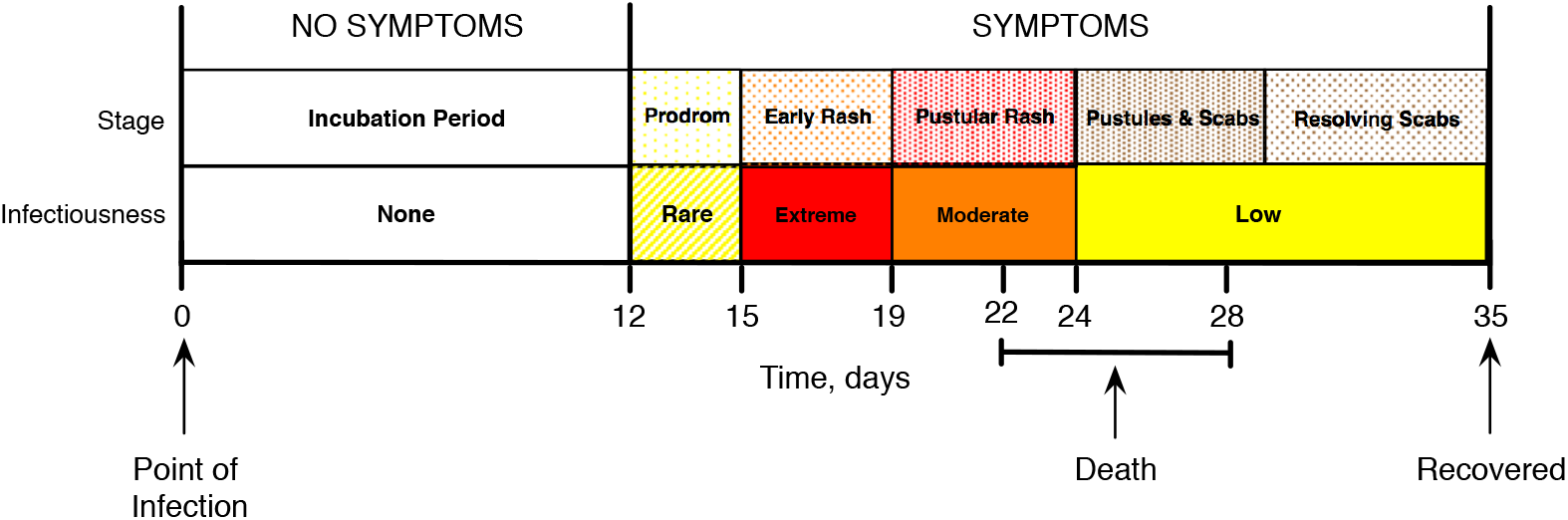
The natural history of smallpox infection. The *prodrom stage* begins with fever but the patient is very rarely contagious. *Early rash* is the most contagious stage, when the rash develops and transforms into bumps. During the *pustular rash* stage bumps become pustules, then turn into scabs during the *pustules and scabs* stage, and finally fall off during the *resolving scabs* stage. The infected person is contagious until the last scab falls off.

There is an *incubation period* during which the infected person has no symptoms and is not contagious. The duration of this stage can vary from 7 to 19 days, but in most cases is about 12 days. The *prodrom* stage begins with the onset of fever, and sometimes includes vomiting and diarrhea. The host is rarely contagious at this stage. The rash appears 2-4 days after the onset of fever. It starts as small red spots on the tongue and in the mouth that grow into sores that break open within 24 hours of their appearance. At this point, a large amount of virus is contained in the mouth and the throat of the infected host, making them extremely contagious. Then the rash spreads rapidly all over the body, and in a few days transforms into bumps filled with thick fluid. This *early rash* stage continues for about 4 days. It is followed by a *pustular rash* stage (average duration 5 days) during which the bumps become pustules. Over the next 5 days (*pustules and scabs* stage), pustules turn into scabs. The scabs fall off during the *resolving scabs* stage (average duration 6 days), often leaving pock marks on the skin. The overall duration of the illness is about 23 days [27]. In fatal cases, the majority of deaths occur on the 10th-16th day from the beginning of symptoms [1, p. 22]. The usual cause of death is severe toxemia, the release of toxins in the blood [1, p. 130] [28, p. 345].

### Data

We accessed original documents in London, England, in the Guildhall Library, the British Library, the Wellcome Library, and the London Metropolitan Archive. We digitized birth, death and population records for London throughout the period over which smallpox was listed as a cause of death (1661-1934). The last smallpox death reported in London was in the week of 24 February 1934.

All data analyzed in this paper are shown in **Fig 2**, and are available both in supplementary material and at the International Infectious Disease Data Archive (IIDDA, http://iidda.mcmaster.ca^1^).

**Fig 2.**
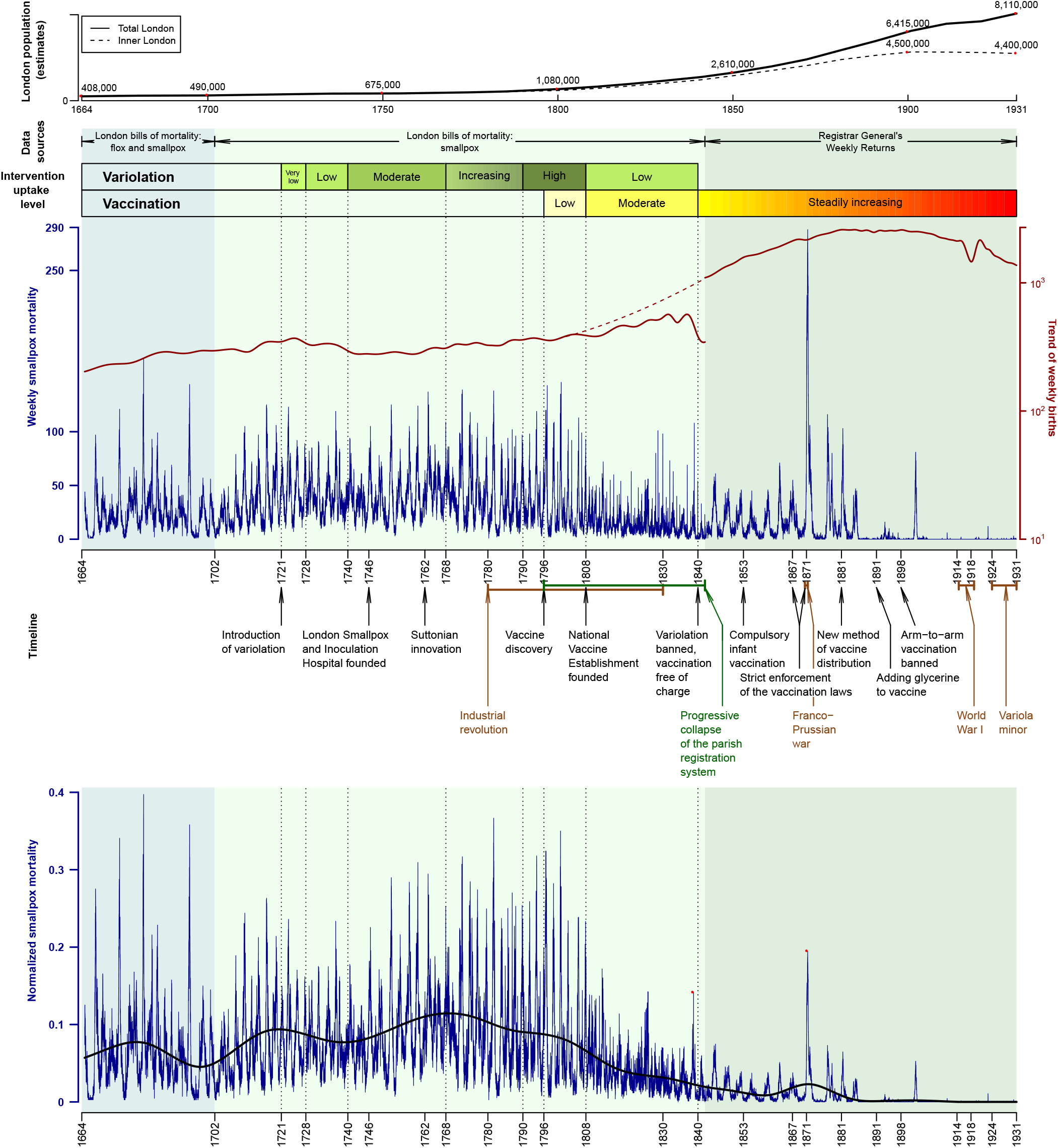
London’s population and weekly smallpox mortality 1664-1930 against the timeline of historical events related to the history of smallpox in England. The top of the graph shows estimates of the London population for inner London (dashed black line) and all of London (solid black line). The top of the main panel (shaded) indicates the data sources. Smallpox mortality data were collected from two sources: i) London Bills of Mortality, where smallpox deaths were recorded from 1664 until 1701 under the name “flox and smallpox” (light blue background colour) and under the name “smallpox” (light yellow background colour) from 1701 until 1841; ii) Registrar General’s Weekly Returns (light grey background colour) used from 1842 until 1931. Intervention uptake levels are shown as coloured bars: yellow green-dark olive for variolation with yellow green indicating the lowest level and dark olive the highest; and yellow-red for vaccination with light yellow indicating the lowest level and red the highest level. Weekly smallpox mortality is presented in dark blue. The trend of weekly births are plotted in dark red. The transition from one registration system to another during 1796–1842 resulted in reduced accuracy. A dashed dark red line shows a fitted birth trend during this period. Below the main panel, we annotate the timeline of historical events related to smallpox history in England: black text indicates events that influenced uptake of control measures; brown text shows events that influenced human behaviour; dark green shows the period when data accuracy was reduced. The **bottom panel** shows a time plot of weekly smallpox mortality normalized by the trend of weekly all-cause mortality (see **Fig S1**). The trend of normalized smallpox mortality (solid black curve) was estimated by Empirical Mode Decomposition. Red dots denote the peaks of the epidemics of 1838 and 1871–1872, the most significant smallpox epidemics of the 19th century.

### London Bills of Mortality

Registration of deaths in England began in 1538 [29, p. 54], but systematic summaries of deaths categorized by cause were not published until later, in the *Bills of Mortality*. The Bills of Mortality included information about baptisms and church burials, categorized by cause of death. The Company of Parish Clerks, who published the bills, collected counts from the individual Anglican parish registers [30,31]. Weekly bills began to be published frequently in 1604 [32], but not without large gaps until 1661. A nearly continuous weekly sequence of bills exists starting^2^ 18 October 1664.

### Reporting of smallpox deaths

Before 1701, smallpox was listed in the Bills of Mortality under the heading *“flox and smallpox”. “Flox”* is an old term for the haemorrhagic type of smallpox [32,33, p. 436]. After 1701, *“smallpox”* was used consistently. Other disease names that occur in the bills and are considered by some to be associated with smallpox are *“flux”* and *“bloody flux”*. Razzell [3] suggested that bloody flux was a name used for haemorrhagic smallpox, and that it was considered a distinct disease [3, p. 104]. However, Creighton [32] described bloody flux as an old name for dysentery and not as something related to smallpox [32, p. 774]. A *Glossary of Archaic Medical Terms* [33] also defines *“bloody flux”* as dysentery and *“flux”* as diarrhea. In any case, mortality from *“bloody flux”* and *“flux”* was negligible compared to smallpox (4,679 deaths in total from “bloody flux” and “flux” compared to 322,219 total smallpox deaths). Consequently, even if they were related to smallpox deaths, they would not significantly influence our findings. We therefore used the sum of *“flox and smallpox”* and *“smallpox”* records and did not include *“bloody flux”* and *“flux”* in our data.

### Transition to the Registrar General’s Weekly Returns

The Bills of Mortality were the only official registration system used in England until the introduction of national records, the Registrar General’s Weekly Returns, in 1837. The necessity for improvement of the methods of data collection became evident at the end of the 18th century. The accuracy of the old system of parish records had been compromised by the rapid growth of London’s population from the beginning of the industrial revolution. Parish clerks simply could not keep up with the increasing flow of information. As a result, the Registrar General’s Office was created with the main purpose of keeping birth and mortality records more complete, and covering all sectors of the population [31].

We used the Registrar General’s Weekly Returns from 1842 onwards.

### Adjustments during the transition period

Throughout the period 1796–1842, the last week of the reported year, which was the first week of December, had an unusually high number of reported deaths. The reason for this appears to be backlog, *i.e.*, that deaths that had occurred previously (but were not counted at the time) were all reported in the first week of December. To address this inaccuracy, we replaced the records showing strikingly high numbers of deaths with the average of the previous and following weeks. Then the difference between the original and the replaced values was uniformly distributed throughout the year to keep the annual number of deaths consistent with the original data.

### Missing weeks

The mortality bills for some weeks have been lost. Fortunately, all gaps are small (typically 1–5 weeks) with the largest gap being 9 weeks. These gaps were replaced with linearly interpolated values to obtain a time series without missing values.

### Consistency checks

Annual summaries of mortality in London were published from 1629 onwards. Initially, these were annual bills of mortality and later they were annual summaries of the Registrar General’s returns. Creighton [32] tabulated annual all-cause mortality (1629-1837) and smallpox mortality (1629-1893). In Fig S2, we compare these annual counts with annual aggregations of our weekly data. The data from the two sources align well for most years. There are significant differences only during 1837-1841, which was the period of transition of control of mortality summaries from the Company of Parish Clerks to the Registrar General for England and Wales.

### London’s population

Decennial censuses of London began in 1801 [34]. We estimated London’s population at earlier times based on Finlay and Shearer [35] for the period before 1700 and Landers [36] for 1700-1800. See Appendix.

## Annotation of data with historical events

Over the 267 years that we study here, London underwent major demographic and social changes, and there were a variety of historical events that may have had substantial impacts on smallpox dynamics. The changes that were most obviously relevant to smallpox dynamics were the introduction of smallpox control measures (variolation and vaccination). Other potentially relevant events include wars and the industrial revolution. We annotate the smallpox time series in **Fig 2** with the major events and developments that we described below.

### Control-free era

The first recorded outbreak of smallpox in England is dated to 12 August 1610 [32, p. 435]. Smallpox became one of the major causes of death in England in the 17th century, exceeding plague, leprosy and syphilis [2,31,36]. The impact of smallpox did not change until preventative measures were introduced at the beginning of the 18th century.

### Early variolation, 1721–1740

Lady Mary Wortley Montagu (an influential writer and poet [37]) is credited with introducing variolation to Great Britain [1–3,17,38]. She had her daughter professionally variolated in London in April 1721. For the next two decades, variolation occurred but was not popular. Only 857 people were variolated in the whole of Great Britain from 1721 to 1727, and only 37 in 1728 [32]. After 1728, precise numbers are unknown but assumed to be low. English medical practitioners implemented variolation very crudely with deep incisions that caused severe symptoms, morbidity, and high mortality of up to 2%.

### Increasingly common variolation, 1740–1808

At the beginning of the 1740s, only the rich could afford variolation. Charitable variolation began with the establishment of the London Smallpox and Inoculation Hospital in 1746.

Until 1762, variolation was usually preceded by four to six weeks of preparation, which included purging, bleeding, and a restricted diet with limited amount of food. Variolation was followed by an isolation period of two or more weeks. Isolated individuals were placed in purpose-built inoculation houses [1,3, p. 255]. The preparation period was shortened after Robert Sutton’s improvement of variolation using light incisions in 1762. Sutton’s new method dramatically decreased the severity of symptoms and death, and reduced the cost. Consequently, it became more common to offer variolation to the poor population free of charge.

Sutton’s variolation technique spread quickly around England and became very popular in rural areas. When a new epidemic appeared to be highly probable, “general variolation” of entire villages and communities was performed. In large towns and cities the situation was quite different [39]. In London, the use of variolation was irregular and attempts to perform “general variolation” were sporadic and rare.

Poor Londoners were variolated only through Smallpox Charities. They performed public variolation in batches, separately for males and females, 8–12 times a year [32, p. 506]. The charities did not admit children under 7 years of age despite the fact that the vast majority of smallpox cases at that time were in children under 3 years of age.

The full extent of variolation in London after the Suttonian innovation is unclear. The figures from London Hospital show that the number of variolated individuals increased dramatically from 29 in 1750 to 653 in 1767 and 1084 in 1768 [32, p. 506]. Based on a variety of historical reports, Razzel concluded that variolation gained considerable popularity in London at the turn of the 19th century [3, p. 72]. We consider uptake of variolation to have been moderate during 1740–1768, after which update increased and reached a maximum during 1790–1808 [3,40].

### Industrial revolution

Beginning in the 1780s, the Industrial Revolution brought thousands of immigrants to London and led to an increase in population density [41–43]. Smallpox was probably always present in London, unlike rural settings, which were relatively smallpox free between epidemics [1,44]. Consequently, whereas most adults in London would have acquired smallpox when young, immigrants from rural areas were more likely to have escaped infection, and therefore contributed continously to the flow of susceptibles into London. The larger susceptible pool might explain the increasingly severe smallpox outbreaks at the end of the 18th century, despite the growing popularity of variolation (**Fig 2**).

### Vaccination

The discovery of a vaccine by Edward Jenner in 1796 was the most important event in the history of smallpox and was directly responsible for the eventual eradication of the pathogen. At first, however, the idea of vaccination was met with skepticism by the scientific and medical community [1,2]. Indeed, Jenner “was advised not to send a record of his observations to the Royal Society, which was prepared to refuse it, but to publish it as a pamphlet; and as a pamphlet it appeared in 1798” [45, p. 62].

Unlike variolation, vaccination came with relatively little risk to the vaccinee, no preparatory period, and much lower cost. Consequently, vaccination was adopted by the public more quickly and widely than variolation ever was [1]. Nevertheless, there were initially many challenges to overcome associated with ineffective vaccine distribution and storage, shortage of cowpox virus, inadequate vaccine efficacy, and religious and philosophical objections. Despite these difficulties, vaccine uptake rose and led to a dramatic decline in smallpox mortality by the end of the 19th century (**Fig 2**).

Unfortunately, quantitative reports on early smallpox vaccine uptake are sorely lacking. Vaccinations were poorly recorded until the end of the 19th century. Available data are incomplete, uncertain and inconsistent. For example, the figures from the *London Smallpox and Inoculation Hospital* show the percentage of vaccinated patients admitted to the hospital increasing steadily from 32% in 1825 to 73% in 1856 [46, p. 114], whereas the *Royal Commission on Vaccination* found that only 25% of newborns were vaccinated by 1820 and about 70% in some parishes by 1840 [31]. Mooney [47] states that during 1854-1856 the percentage of vaccinated infants might have ranged from 28% to 81% based on one source, but that another source indicates that infant vaccination rates for London during the period 1845-1890 were much lower than the national average and never increased above 500 per 1,000 live births (*i.e.*, 50%). There appear to be no surviving records concerning vaccinations of older age groups during this period; however, it is hypothesized that many adults escaped vaccination [46]. In **Fig 2**, we indicate the qualitative pattern of change in vaccine uptake.

### Better access and increasingly strong legislation concerning vaccination

State involvement in the control of smallpox in England began with the foundation of the *National Vaccine Establishment* in 1808. It provided free vaccination at its London stations and distributed vaccine to other parts of England [48]. Around this time, the *London Smallpox and Inoculation Hospital* ceased variolation and began vaccination in greater numbers. Smallpox outbreaks over the next decade were very mild, which was believed to be due to the growing popularity of vaccination.

A strikingly large smallpox epidemic occurred in London in 1837–1838, and exploded into a European wide pandemic [2]. The authorities in England realized that some radical measures had to be taken, which led to the first *Vaccination Act of 1840*, providing vaccination free of charge and banning variolation. It was followed by the *Vaccination Act of 1853*, which made vaccination of every child during the first four months of life compulsory. The *Vaccination Act of 1867* introduced penalties for not complying with compulsory vaccination.

The Franco-Prussian war, which began in late July 1870, is believed to have initiated the worst pandemic of smallpox in all of Europe in the 19th century. It resulted in at least 500,000 deaths. England alone lost more than 40,000 people. Thanks to compulsory vaccination, fatality rates in England were three times lower than in Prussia, Austria and Belgium [2, pp. 87–91]. The immediate response of the English government to this devastating pandemic was the *Vaccination Act of 1871*, which enforced very strict control (through the courts) of the implementation of the previous Acts.

In the second half of the 19th century, many vaccine-related challenges were resolved. Arm-to-arm vaccination^3^ was the main method of vaccine distribution in the beginning of the century. It was dangerous because it could transmit other diseases such as syphilis, and was consequently outlawed in 1898 [2]. It was replaced by the new technique of passing cowpox from cow to cow. The new method of vaccine distribution was first introduced in Naples, Italy in 1843 [49,50]. However it arrived in England only in 1881. Another important discovery was made in 1891 by Monckton Copeman [50], who demonstrated that adding glycerine to smallpox vaccine reduces bacterial contamination, making it more efficacious and reliable.

### Eradication in the 20th century

The last large outbreak of *Variola major* occurred in London in 1901–1902 (**Fig 2**) and was probably seeded from another country [46]. After 1902, only very small outbreaks occurred with very low incidence and very few deaths.

In 1967, the World Health Organization launched its global smallpox eradication campaign, and by 1980, smallpox was certified as the first infectious disease to be eradicated by human efforts [14].

## Methods

We used a variety of methods to elucidate the patterns in the London smallpox time series. Spectral analyses and a visualization of seasonality allow us to reveal important characteristics of the data that cannot be gleaned by inspection of the raw time plot (**Fig 2**).

### Normalization

To control for changes in population size, city boundaries and data collection methods, we normalized the smallpox data by the trend of the weekly all-cause mortality in London. To calculate the trend, we used **Empirical Mode Decomposition (EMD)**, which is designed to identify trends in nonlinear and highly non-stationary time series [51–53]. EMD was developed to overcome the drawbacks of moving averages, other linear filters, or linear regression, which often perform poorly on non-stationary data. EMD decomposes a signal into several components with a well defined instantaneous frequency via “intrinsic mode functions” (IMF). IMFs are basically zero-mean oscillation modes present in the data: the first IMF captures the high frequency (shorter period) oscillations while all subsequent IMFs have lower average frequency (longer period). Each IMF is extracted recursively starting from the original time series until there are no more oscillations in the residue. The last residual component of this process can be considered an estimate of the trend [52].

### Spectral analysis

We used spectral analyses to identify the strongest periodicities in the smallpox time series.

### Classical power spectrum

We computed the standard power spectral density [54–57] of the entire (normalized and square root transformed) smallpox time series to obtain a global estimate of its frequency content. We employed a standard modified Daniell smoother using the spec.pgram function in R [58].

### Wavelet spectrum

Because infectious disease time series are typically highly non-stationary, wavelet analysis has become increasingly popular in epidemiological research [59–64]. We computed a wavelet transform [65,66] of the (normalized and square root transformed) smallpox time series in order to examine how smallpox periodicities changed over the course of the three centuries.

A wavelet transform is computed with respect to a basic shape function, the *analyzing wavelet*, which has a *scale parameter* that controls its width. Narrower (wider) scales correspond to higher (lower) frequency modes in the time series. At any given time and scale, a stronger correlation between the analyzing wavelet and the signal yields a larger value of the wavelet transform. We obtain a complete (two-dimensional) time-frequency representation of the data by convolving the analyzing wavelet—at each scale—with the original time series. We used the *Morlet* wavelet [66] as the analyzing wavelet to obtain the wavelet transform of the smallpox time series.

The standard computation of the wavelet spectrum requires that the number of points in the time series be a power of 2. Consequently, we pad the ends of the time series with zeros to bring the number of time points in the data to the nearest power of 2. The resulting artificial discontinuity where the zero-padding begins reduces the accuracy of the wavelet transform at the ends of the time series. Regions of lower accuracy are identified by the “cone of influence”. Data outside this cone should be interpreted with caution. 95% confidence regions are computed based on 1,000 Markov bootstrapped time series [60, 61].

### Seasonality

An indication of underlying seasonality in a time series is the occurrence of a spectral peak at a period of one year. However, this crude measure suppresses the detailed seasonal pattern and, in particular, does not reveal the times of year when peaks or troughs occur. Following Tien *et al* [67], we visualized the evolving seasonal pattern of smallpox dynamics with a heat map in the time-of-year *vs*. year plane.

## Results

The time plot of the raw data (**Fig 2**) displays substantial changes in the structure, amplitude, and frequency of smallpox epidemics over time. However, some of the apparent changes are misleading because they do not account for population growth and inconsistency of data sources. For example, the epidemic of 1871 appears to be the largest, but it was not the largest relative to population size. In contrast, the epidemic of 1838, which is frequently mentioned in the literature [2,32], is not easily identifiable in the raw data, likely because it occurred just at the time of transition between the Bills of Mortality and the Registrar General’s Returns.

### Normalization

**Fig S1** shows the trend of weekly all-cause mortality in London (1661–1930) computed with EMD. The bottom panel of **Fig 2** shows the weekly London smallpox time series normalized by the all-cause trend.

The normalized time series provides a more consistent and informative representation of smallpox dynamics in London. For example, the epidemic of 1838 is now easily identifiable, and the epidemic of 1871–1872 is much less extreme in magnitude compared with other epidemics of the 19th century.

From the earliest times in the series, there were large, recurrent epidemics. More common variolation after 1770 is correlated with stricter regularity of epidemics, while the introduction of vaccination is associated with a dramatic reduction in the amplitude of epidemics. During the period when variolation and vaccination were both in use (1796–1840), the data appear to be noisier, and outbreaks occurred more frequently. After variolation was banned in 1840, inter-epidemic intervals increased and epidemic peak heights declined, except for three large epidemics in 1871, 1876 and 1902.

The bottom panel of Fig 2 also shows the general trend of smallpox mortality over the centuries (computed with EMD). This trend indicates that *per capita* smallpox mortality increased steadily from 1664 until approximately 1770, after which there was a gradual decline until its complete elimination. The begining of the decline is coincident with the growing popularity of variolation. London’s population grew by an order of magnitude over the subsequent years from 1796 when the vaccine was discovered until 1930; yet smallpox mortality in the city fell to negligible levels as vaccine uptake rose.

### Spectral analysis

#### Classical power spectrum

**Fig 3** shows the period periodogram (power spectrum as a function of period) for the full smallpox time series. A strong peak at one year suggests underlying seasonality of epidemics. Other peaks (near 2.2, 2.4, 3, 5.1 and 6 years) suggest more complex dynamical patterns.

**Fig 3.**
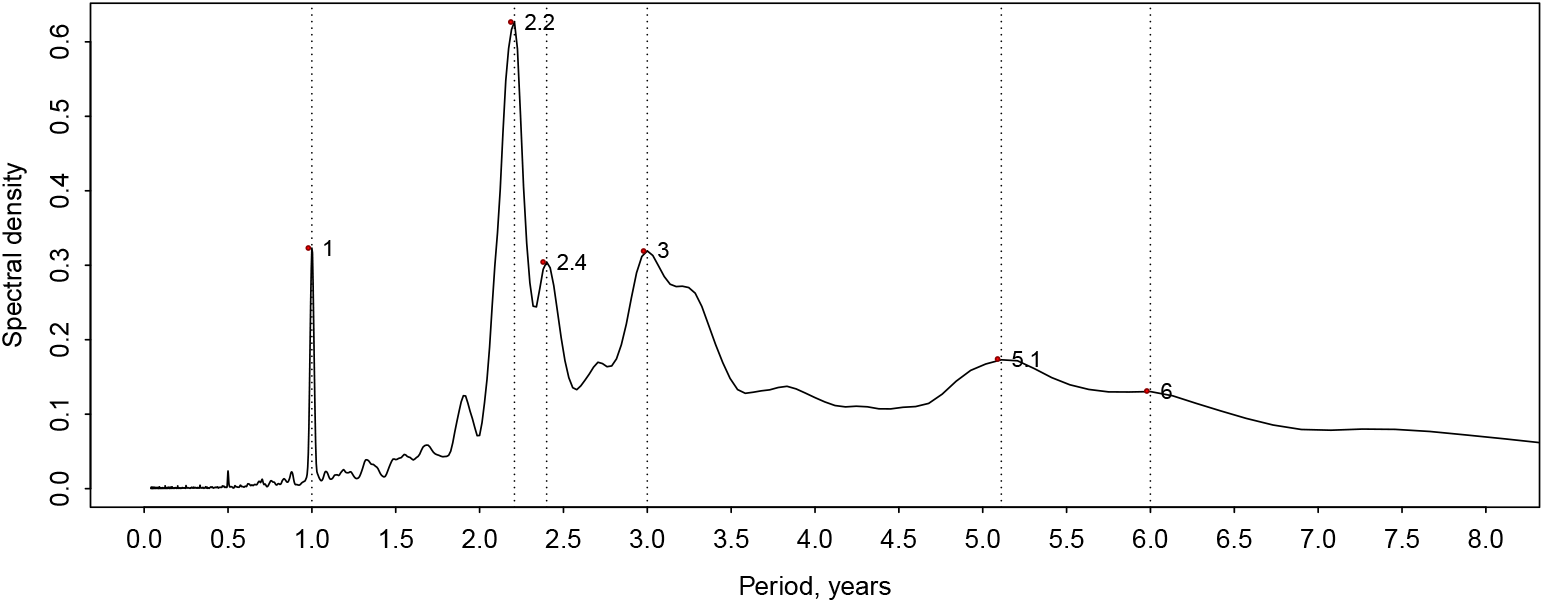
Classical Fourier power spectrum of the normalized weekly smallpox mortality time series for London, England, 1664-1930. Before computing the power spectrum the time series was detrended and square root transformed [54]. The spectrum was smoothed using a modified Daniell window (weighted moving average) [54].

#### Wavelet spectrum

The bottom panel of **Fig 4** shows the wavelet transform of the London smallpox time series. Colours indicate strength of signal at given periods (blue meaning weak and red meaning strong). The cone of influence and 95% confidence contours are shown in black.

**Fig 4.**
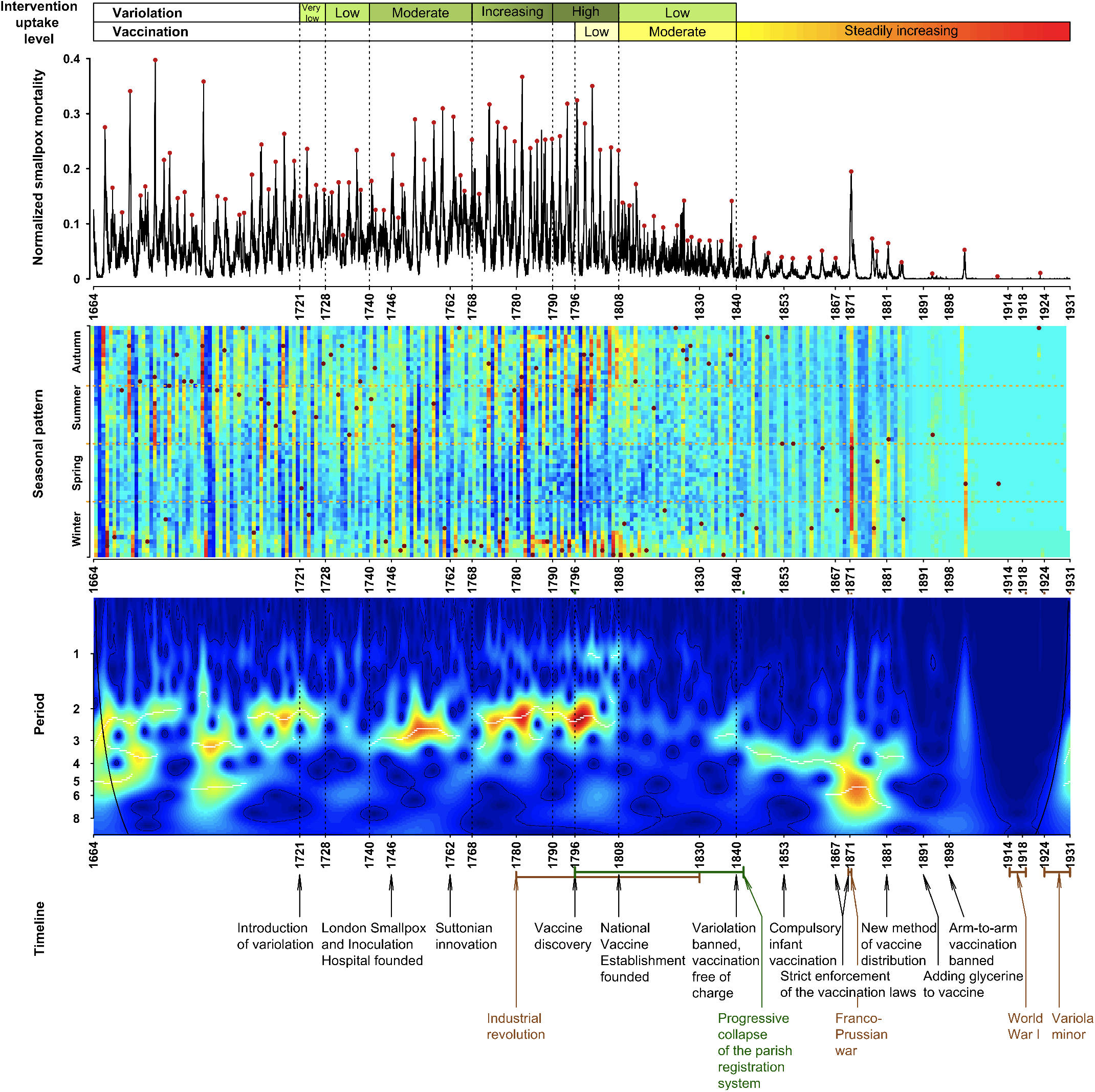
Spectral and seasonal structure of smallpox dynamics in London, England (1664-1930). **Top panel**: Weekly smallpox mortality normalized by the trend of all-cause mortality together with variolation and vaccination uptake levels, with red dots highlighting the major peaks in the time series (identified by visual inspection). **Middle panel**: Seasonality of smallpox epidemics. The normalized smallpox time series was detrended and square root-transformed before constructing the heat map. Dark red dots indicate the week of each year with the highest value of normalized smallpox mortality (as in the time series in the top panel). Zeros are not necessarily displayed with dark blue, because the detrending shifts the end of the time series up. **Bottom panel**: Wavelet transform of the normalized weekly smallpox mortality time series (square root-transformed and normalized to unit variance). The white curves show the local maxima of the wavelet power (squared modulus of wavelet coefficients [61, p. 291] at each time). The colours of the wavelet diagram vary from dark blue for low power to dark red for high power. The thin black line indicates the 95% confidence region, estimated from 1000 bootstrapped time series generated by the method of [61, pp. 292-293]. Below the “cone of influence” [61,66], the calculation of wavelet power is less accurate because it includes edges of the time series that have been zero-padded to make the length of the series a power of 2. The wavelet spectrum was computed using MATLAB code kindly provided by Bernard Cazelles. As in **Fig 2**, the graph is annotated with the timeline of historical events related to smallpox history in England.

The white curves in bottom panel of **Fig 4** highlight the period with greatest power at each timepoint. From 1664 until 1700 the dominant period was 3-4 years. Around 1705, it shifted to 2-3 years. After the introduction of variolation, wavelet power weakens overall and the dominant period lengthens. From around 1740 to 1770, a 3-year mode dominates. Between 1770 and 1810, both one year and 2–3 year periods are strong. After 1820, annual power is much less clear, and after 1840 the major period smoothly drifted into a longer cycle of 3–4, and then 4–8 years.

### Seasonality

The middle panel of **Fig 4** shows the seasonal structure of smallpox mortality in London. Until 1740, the maximum number of smallpox deaths occurred in the summer or autumn. Then, until about 1770, outbreak peak times shifted to winter. After 1770, the majority of deaths occurred in the autumn and winter. Epidemic patterns became much less regular and less seasonal during the period 1808–1840. From the beginning of the data set until 1840, smallpox mortality was lowest in the spring. After 1840, as vaccination levels gradually increased, epidemics became strictly regular, with the majority of deaths occurring in winter and spring. After the exceptionally large epidemic of 1871, smallpox epidemics became extremely irregular and noisy.

## Discussion

We have digitized and analyzed what is, to our knowledge, the longest existing weekly time series of infectious disease mortality. The timespan of the data, from 1664 to 1930, covers an extraordinary period in London, England, during which smallpox changed from a terrifying and unavoidable danger to an easily preventable infection.

Previous studies of smallpox dynamics in London were based on annual mortality records [12,44,68], which preclude the detection of seasonal patterns. Our spectral analysis revealed the presence of annual periodicity through the smallpox time series, and changes in the strength of annual power are correlated with changes in control practices. The previous work [12,44,68] attributed changes in inter-epidemic intervals to changes in birth rates and nutrition; while these factors are likely to have influenced smallpox dynamics in London, the correlations we have highlighted between the uptake of preventative measures (and related legislation) and the observed changes in epidemic patterns, strongly suggest that transitions in epidemic patterns were primarily driven by control interventions.

Seasonal forcing of infectious disease transmission can stimulate persistence of complex epidemic cycles [69] and chaos [70]. The mechanistic origin of seasonal forcing may [71] or may not [72] be easy to determine. In the case of smallpox, previous work [1,12,73] found that in temperate climates the majority of smallpox cases occurred in the winter and spring, whereas in tropical climates the seasonality was less pronounced. The general conclusion was that smallpox incidence always increases when the weather is cool and dry; this belief influenced the planning of the eradication campaign in India and seemed to help to improve its efficiency [1, pp. 179–181]. Previous smallpox seasonality studies were mainly based on data from the 19th and the 20th centuries, when preventative measures were already common [1,73]. Our data set is of a particular interest in this respect since it includes a period when only naturally acquired smallpox immunity existed. Consequently, we were able to compare early and later periods (which displayed a shift from summer to winter epidemics) and comment on the impact that intensive preventive measures appeared to have on smallpox seasonality (much greater regularity until smallpox deaths were mostly eliminated).

Better control is naturally expected to lead to less death. However, how interventions influence the frequency structure and seasonality of epidemic time series over decades and centuries is much more subtle [56,74,75]. While preliminary work has been promising [76], careful estimation [77,78] and analysis [75,79] of patterns of seasonal forcing will be required in order to use mechanistic models reliably to explain [64] the observed transitions in smallpox transmission dynamics.

## Appendix: Population of London (1550—1931)

Finlay and Shearer [35] estimated the total population of London for every half century from 1500 until 1700 based on information about the population north and south of the river Thames, taking into account inaccuracies in data collection. Their book [35] also contains data for other years, which was not included in their summary table. We included those additional data in our study as well. Note that Finlay and Shearer estimate for 1650 appears to be inaccurate due to an arithmetic error (it should be ≈400,000 instead of 375,000 according to their own figures). Population data for 1700–1800 were taken from Landers [36], who estimated the population for every decade from 1730 until 1800. Census data, which is available for each decade since 1801, were the source of remaining values [34]. The resulting data set of population estimates is plotted in the top panel of **Fig 2**.

**Table 1.** Population of London, England (1550-1931).

| Year | Total London | Outer London | Inner London | Source |
| --- | --- | --- | --- | --- |
| 1550 | 120,000 | 120,000 |  | [35] |
| 1560 | 140,800 | 140,800 |  | Estimated<sup>1</sup> |
| 1580 | 180,400 | 180,400 |  | Estimated<sup>1</sup> |
| 1600 | 200,000 | 200,000 |  | [35] |
| 1620 | 297,000 | 297,000 |  | Estimated<sup>1</sup> |
| 1640 | 390,500 | 390,500 |  | Estimated<sup>1</sup> |
| 1650 | 391,450 | 391,450 |  | Estimated<sup>1</sup> |
| 1660 | 392,400 | 392,400 |  | Estimated<sup>1</sup> |
| 1680 | 468,600 | 468,600 |  | Estimated<sup>1</sup> |
| 1700 | 490,000 | 490,000 |  | [35] |
| 1735 | 660,000 | 660,000 |  | [36] |
| 1745 | 670,000 | 670,000 |  | [36] |
| 1750 | 675,000 | 675,000 |  | [35] |
| 1755 | 680,000 | 680,000 |  | [36] |
| 1765 | 730,000 | 730,000 |  | [36] |
| 1775 | 780,000 | 780,000 |  | [36] |
| 1785 | 859,234 | 826,502 |  | [36] |
| 1795 | 1,007,703 | 909,507 |  | [36] |
| 1801 | 1,096,784 | 959,310 | 137,474 | [34] |
| 1811 | 1,303,564 | 1,139,355 | 164,209 | [34] |
| 1821 | 1,573,210 | 1,379,543 | 193,667 | [34] |
| 1831 | 1,878,229 | 1,655,582 | 222,647 | [34] |
| 1841 | 2,207,653 | 1,949,277 | 258,376 | [34] |
| 1851 | 2,651,939 | 2,363,341 | 288,598 | [34] |
| 1861 | 3,188,485 | 2,808,494 | 379,991 | [34] |
| 1871 | 3,840,595 | 3,261,396 | 579,199 | [34] |
| 1881 | 4,713,441 | 3,830,297 | 883,144 | [34] |
| 1891 | 5,571,968 | 4,227,954 | 1,344,014 | [34] |
| 1901 | 6,506,889 | 4,536,267 | 1,970,622 | [34] |
| 1911 | 7,160,441 | 4,521,685 | 2,638,756 | [34] |
| 1921 | 7,386,755 | 4,484,523 | 2,902,232 | [34] |
| 1931 | 8,110,358 | 4,397,003 | 3,713,355 | [34] |

## Acknowledgments

The data were photographed and/or entered primarily by Kelly Hancock, Claire Lees, James McDonald, Laxmi Pandit, and David Richardson. Valerie Hart facilitated work at the Guildhall Library in London. We were supported by the Natural Sciences and Engineering Research Council of Canada (O.K. by an NSERC Postgraduate Scholarship and D.E. by an NSERC Discovery grant). Early stages of this research were supported by a J. S. McDonnell Foundation Research Award to D.E. (https://www.jsmf.org/grants/d.php?id=2006014). We thank all members of the Mathematical Biology Group at McMaster University for helpful comments and discussions.

## SUPPLEMENTARY FIGURES

**Fig S1.**
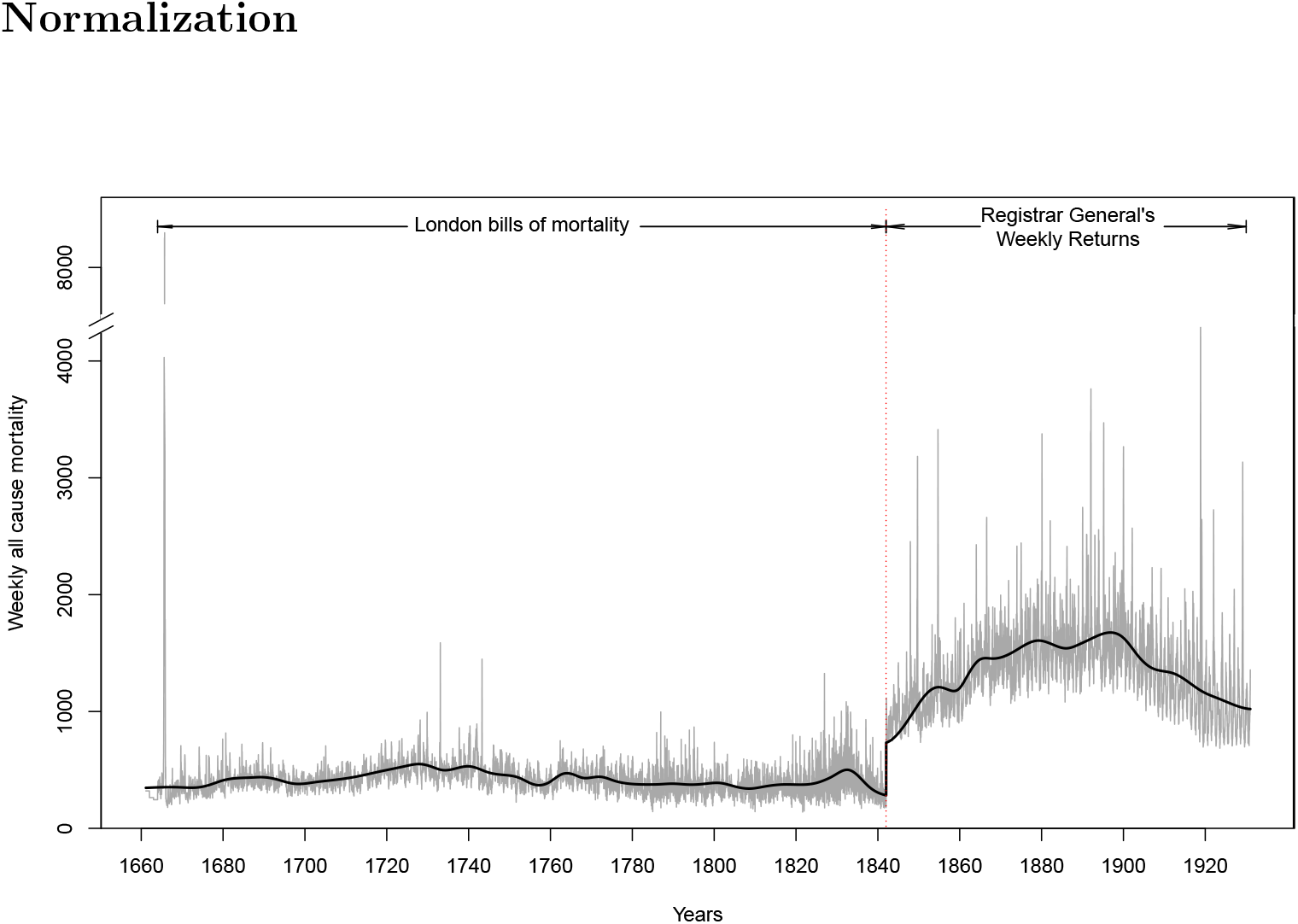
Weekly all-cause mortality and its trend; London, England, 1661–1930. The trend was estimated by Empirical Mode Decomposition applied separately to the periods 1661–1842 and 1842–1930, which correspond to different data sources. The highest peak in all-cause mortality occurred during the Great Plague of London in 1665, which killed more than 8,000 people in the most severe week.

**Fig S2.**
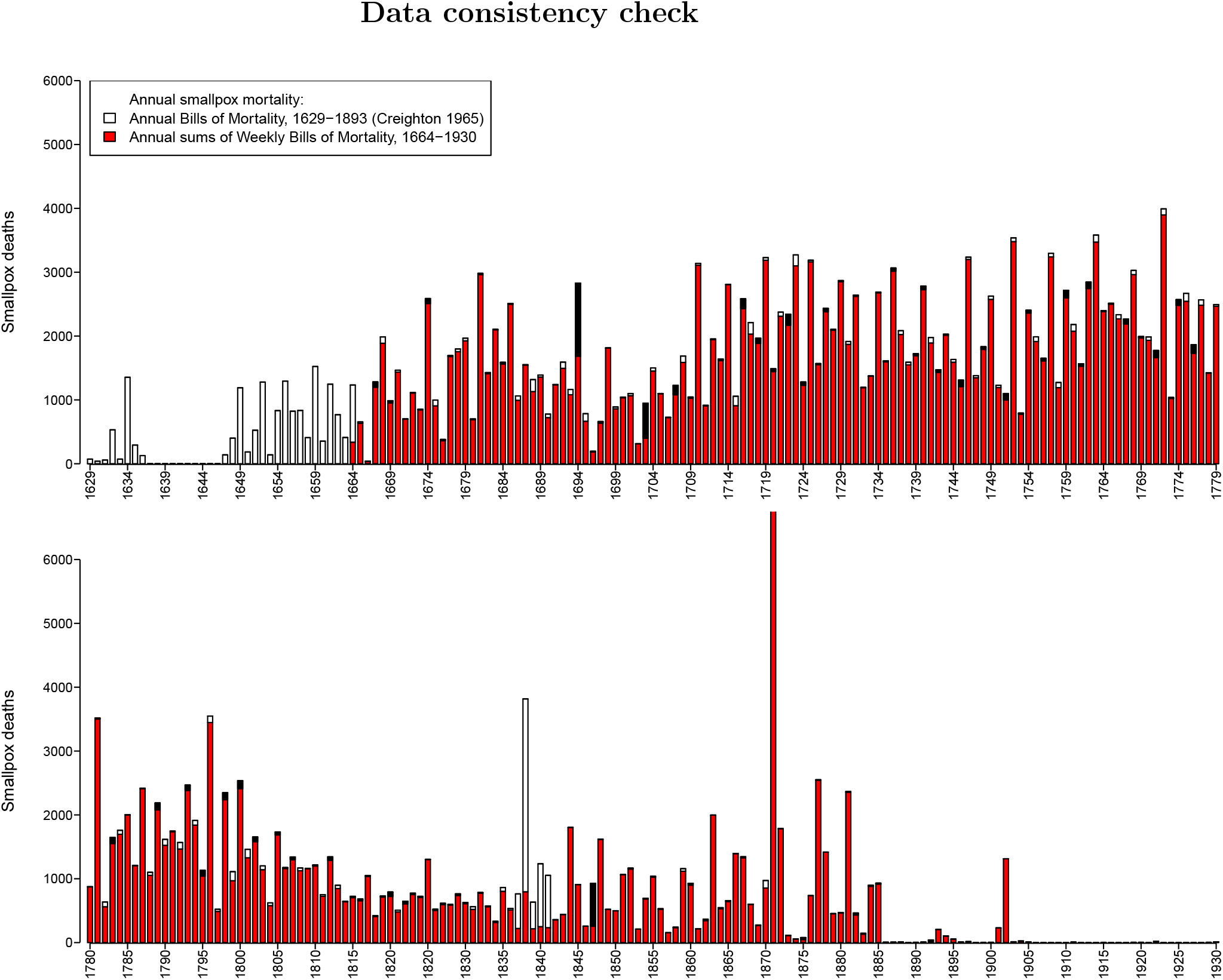
Annual smallpox mortality data in London, England: 1629-1779 (top panel) and 1780-1930 (bottom panel). The data were gathered from two sources: the Annual Bills of Mortality tabulated by Creighton [32] and annual sums of our weekly counts from the weekly Bills of Mortality and the Registrar General’s Weekly Returns. Differences between the two data sets are indicated with white stacked bars if values in the Annual Bills are larger, and black stacked bars if corresponding annual sums from the weekly bills are larger. Note that no smallpox mortality data for the period 1637-1646 were available [32].

## Footnotes

1 Please note that at the time of submission of this paper, the IIDDA web site is down while major restructuring is taking place. We hope this will be completed before the paper is published.

2 Until 1752, New Years Day was 25 March in England. However, we always indicate years following the modern practice of defining New Years Day to be 1 January.

3 Arm-to-arm vaccination was the method of vaccine distribution, which involved vaccine transfer from the infectious pustule of vaccinated individual to a non-vaccinated individual, and so on [1].

